# Long antiparallel open reading frames are unlikely to be encoding essential proteins in prokaryotic genomes

**DOI:** 10.1101/724807

**Authors:** Denis Moshensky, Andrei Alexeevski

## Abstract

The origin and evolution of genes that have common base pairs (overlapping genes) are of particular interest due to their influencing each other. Especially intriguing are gene pairs with long overlaps. In prokaryotes, co-directional overlaps longer than 60 bp were shown to be nonexistent except for some instances. A few antiparallel prokaryotic genes with long overlaps were described in the literature. We have analyzed putative long antiparallel overlapping genes to determine whether open reading frames (ORFs) located opposite to genes (antiparallel ORFs) can be protein-coding genes.

We have confirmed that long antiparallel ORFs (AORFs) are observed reliably to be more frequent than expected. There are 10 472 000 AORFs in 929 analyzed genomes with overlap length more than 180 bp. Stop codons on the opposite to the coding strand are avoided in 2 898 cases with Benjamini-Hochberg threshold 0.01.

Using Ka/Ks ratio calculations, we have revealed that long AORFs do not affect the type of selection acting on genes in a vast majority of cases. This observation indicates that long AORFs translations commonly are not under negative selection.

The demonstrative example is 282 longer than 1 800 bp AORFs found opposite to extremely conserved *dnaK* genes. Translations of these AORFs were annotated “glutamate dehydrogenases” and were included into Pfam database as third protein family of glutamate dehydrogenases, PF10712. Ka/Ks analysis has demonstrated that if these translations correspond to proteins, they are not subjected by negative selection while *dnaK* genes are under strong stabilizing selection. Moreover, we have found other arguments against the hypothesis that these AORFs encode essential proteins, proteins indispensable for cellular machinery.

However, some AORFs, in particular, *dnaK* related, have been found slightly resisting to synonymous changes in genes. It indicates the possibility of their translation. We speculate that translations of certain AORFs might have a functional role other than encoding essential proteins.

Essential genes are unlikely to be encoded by AORFs in prokaryotic genomes. Nevertheless, some AORFs might have biological significance associated with their translations.

**Author summary:** Genes that have common base pairs are called overlapping genes. We have examined the most intriguing case: if gene pairs encoded on opposite DNA strands exist in prokaryotes. An intersection length threshold 180 bp has been used. A few such pairs of genes were experimentally confirmed.

We have detected all long antiparallel ORFs in 929 prokaryotic genomes and have found that the number of open reading frames, located opposite to annotated genes, is much more than expected according to statistical model. We have developed a measure of stop codon avoidance on the opposite strand. The lengths of found antiparallel ORFs with stop codon avoidance are typical for prokaryotic genes.

Comparative genomics analysis shows that long antiparallel ORFs (AORFs) are unlikely to be essential protein-coding genes. We have analyzed distributions of features typical for essential proteins among formal translations of all long AORFs: prevalence of negative selection, non-uniformity of a conserved positions distribution in a multiple alignment of homologous proteins, the character of homologs distribution in phylogenetic tree of prokaryotes. All of them have not been observed for the majority of long AORFs. Particularly, the same results have been obtained for some experimentally confirmed AOGs.

Thus, pairs of antiparallel overlapping essential genes are unlikely to exist. On the other hand, some antiparallel ORFs affect the evolution of genes opposite that they are located. Consequently, translations of some antiparallel ORFs might have yet unknown biological significance.

## Introduction

Genes that have common base pairs are of interest due to their origin and evolution of overlapping parts [1,2]. In most cases, overlaps of genes consist of a couple of base pairs. Gene pairs that have long overlaps are common for viruses [3–8]. Some authors explain it with the strong limitation of a viral genome length. Overlapping genes were studied by computational methods both in prokaryotic and eukaryotic genomes. However, a few cases of long overlaps were reliably confirmed by experimental studies.

According to Pallejà et al. [9], all studied 715 co-directional gene overlaps longer than 60 bp in prokaryotes seem non-existing. We focused on prokaryotic overlapping genes located on opposite DNA strands, which are called antiparallel overlapping genes (AOGs). Such genes must be translated from different mRNA. Thus they have different promoters and regulatory signals.

To the best of our knowledge, there are experimental works claiming certain AORFs translation in *Escherichia coli* [2,10,11], *Streptomyces coelicolor* [12] and *Pseudomonas fluorescens* [13,14].

Antiparallel overlapping genes *dmdR1* and *adm* were detected in *S. coelicolor* [12]. The *adm* gene is located almost entirely in complement strand of *dmdR1*, having 16 extra nucleotides at 3’ end. The *adm* frame is −2 relative to +1 frame of *dmdR1*. The length of overlap is 686 bp. The *dmdR1* gene encodes a well-characterized conserved major regulator of iron metabolism. Both *dmdR1* and *adm* products were confirmed by western blot analysis using specific antibodies against both proteins. The authors showed that *adm* knockout, retaining *DmdR1* amino acid sequence unchanged, stimulates sporulation and produces much more pigmented antibiotics in specific media then the wild type and other mutants. They concluded that *adm* gene product may act as a negative regulator of antibiotic biosynthesis.

About 1000 bp antiparallel overlap of genes *Pfl_0939* and *iiv14* was detected in *P. fluorescens* Pf0-1 [13]. These AOGs are in −2 frame relative to each other. Protein products were detected for both genes. These proteins seem to affect soil colonization. However, the particular way they do it was not highlighted.

Ten cases of genes overlap were detected by mass-spectrometry in *P. fluorescens* [14], nine of them are AOGs, and the rest appears as co-directional overlap. Transcription of both AOGs was confirmed by reverse transcriptase PCR in all nine cases. The lengths of predicted (novel) proteins are from 42 amino acids to 530 amino acids. The biological role of these proteins remains to be elucidated.

Long (more than 1500 bp) AORF was found [15–17] opposite to the oomycete *Achlya klebsiana* gene of extremely conserved chaperone DnaK (from HSP70 famiily) [18,19] that exists in virtually all living organisms. The translation of that ORF was claimed to have glutamate dehydrogenase (GDH) activity. Analogous long AORFs to the *dnaK* gene were found in other bacteria and eukaryotes [20–23]. Translations of found long AORFs opposite to *dnaK* genes initiated Pfam [24] family NAD-GH (PF10712) of GDHs. All observed long AORFs opposite to *dnaK* gene are in frame -1. Despite the GDH activity of these AORFs translations was placed in doubt [25], they could be translated. The pair *dnaK* and its AORF was suggested as an ancestor of two aminoacyl tRNA synthetases classes [26–29]. Should this dubious hypothesis be the truth, there were AOGs in ancient organisms.

Probably, there is not enough reason to exclude the possibility of long AORFs encoding proteins in prokaryotes. Statistical analysis shows that the number of long ORFs is significantly bigger than expected [30,31]. Unannotated AOGs existence could be the explanation.

Mutations within gene overlaps are constrained [32–34]. Overlap lengths distributions significantly vary with different frames for putative AOGs with overlap size 15-98 bp [35]. It was suggested that an amino acid substitution only in one gene is more likely than amino acid substitutions in both AOGs. AOGs in −2 frame were assumed to be the rarest because, in this frame, the mutation leading to amino acid substitution in one gene results into an amino acid substitution in the antiparallel gene. For two other frames, there are positions in which mutations lead to amino acid substitutions only in one gene but are synonymous for the antiparallel gene. The evolution simulation of a gene under purifying selection [36] shows that its AORF in the frame −2 looks like a protein coding gene under negative selection because third codons positions of the gene and the AORF are opposite to each other – most synonymous mutations in the gene are synonymous in the AORF. For two other frames, if an AORF translation is not under negative selection, it looks like a gene under positive selection.

Several computational methods to predict whether AORF is functional were proposed. One is based on the identification of codon usage bias both in gene and AORF that is in favor of their functionality [37]. Others are based on the special model to calculate Ka/Ks ratio for AORF translation [38,39]. The first model produces false-positives in a large number of cases for AORFs, especially in the frame −2, because codon usage bias in a gene leads to some bias for AORF too. The second model should be used if no selection detected for both gene and AORF: if negative selection detected for one sequence the second sequence is likely not to be under any selection [36].

In this work, we have fetched all AORFs, overlapping with annotated genes by at least 180 bp, from 929 prokaryotic genomes. To define which of them could be protein-coding genes, the conditional probabilities of accident AORFs located opposite functional genes (P-value of stop codon avoidance in AORF) have been calculated. Using Ka/Ks ratio we have identified that the selection affecting genes with long AORFs does not differ from the selection affecting genes without long AORFs. Some genes with long AORFs have been found to be more conservative than other genes both in synonymous and non-synonymous sites. This might be due to slight restrictions on long AORFs sequences variability. Essential proteins (proteins that are indispensable for cellular life [40]) are unlikely to be encoded by long AORFs. However, AORFs translations might play some role for organisms.

## Methods

### Analyzed genomes

We selected 929 prokaryotic genomes (S1 Table) from NCBI RefSeq Chromosome database which we used as input to our analysis. They represent 895 unique species from 518 genera.

### Antiparallel open reading frames search procedure

In this work, a nucleotide sequence between two stop codons in the same reading frame is considered as ORF. The search procedure applied in this work consists of following steps. Annotated genes not shorter 180 bp are selected. Both 3’- and 5’-ends of each chosen gene nucleotide sequence are extended by 180 bp according to a chromosome sequence. The search for all ORFs is conducted in reverse strand of extended genes sequences – found ORFs are AORFs. AORFs overlapped with an annotated gene less than 180 bp are excluded. For each remained AORF a frame is determined (Table 1) and P-value of a stop codon avoidance is calculated.

**Table 1.**
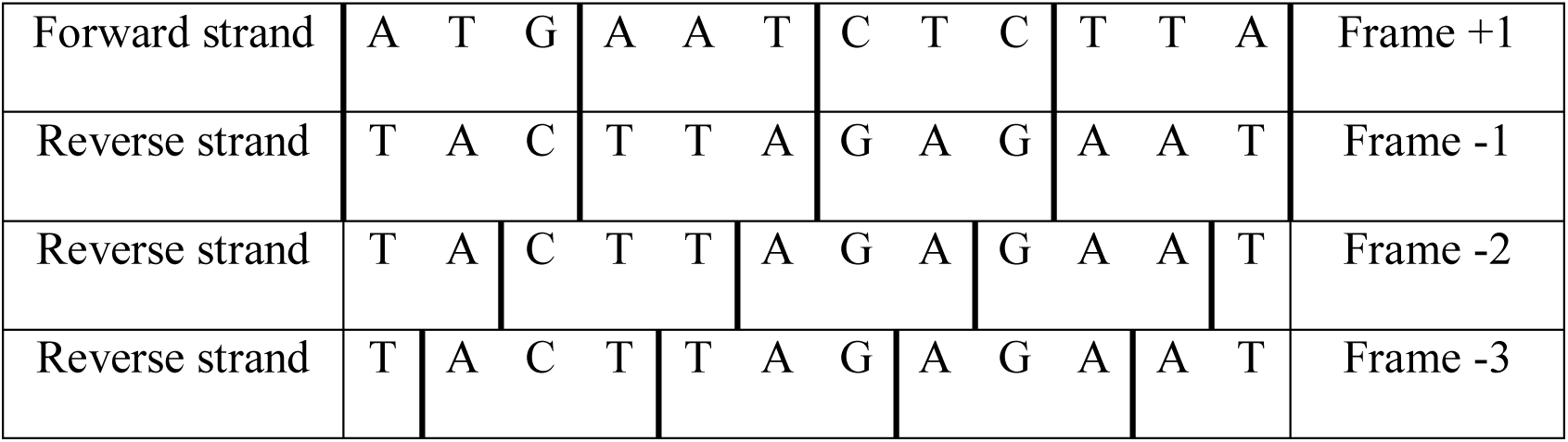
Reading frame position for different ORFs.

### P-value of stop codon avoidance

P-value calculation is based on the assumption that synonymous sites mutate with constant probabilities in all genes for an organism. For a particular genome, these probabilities are taken from codon usage tables. Formulas for P-value calculation vary for AORFs in different frames. In the frame −1, an AORF stop codons TAA, TAG or TGA are located oppositely to codons TTA, CTA or TCA in gene sequence. TTA and CTA encode leucine and TCA encodes serine. The probability of a stop codon absence in an AORF in the frame −1 is calculated by formula 1.

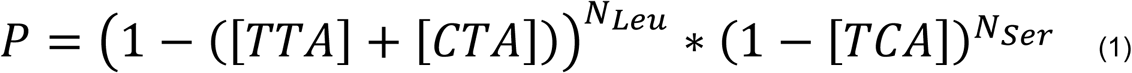

N_Leu_ is the number of leucine encoded in the part of gene overlapped with the AORF, N_Ser_ - the same for serine, [TTA] – frequency of leucine codon TTA for an analyzed organism (the same for CTA and TCA).

A stop codon of an AORF in −2 or −3 frame is located opposite to a pair of codons forming triplets TTA, CTA and TCA. For example, the sequence TCA for AORF in −2 frame will be composed in this manner: NNT|CAN (N – any nucleotide) and for −3 frame NTC|ANN. Consequently, there are pairs of amino acids giving stop codon on the complement strand in a certain frame. For example, the coding sequence of dipeptide Ser-Gln (SQ) contains stop codon TCA on complement strand in −2 frame, if codons TCT or AGT are used for serine and CAA or CAG for glutamine. SQ dipeptide’s coding sequence gives this stop codon with probability equal to ([TCT] + [AGT])*([CAA] + [CAG]). Other stop codons cannot be located opposite to SQ dipeptide. The probability is calculated by formula 2.

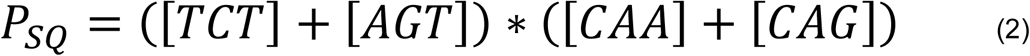

Analogous equations for other dipeptides and for −3 frame are presented in S2 Table. Further, these probabilities are marked as P_ij_, where i,j are numbers of amino acids from 1 to 20.

P-value of AORF in −2 or −3 frame is calculated by formula 3, where n_ij_ is the number of dipeptides ij in gene’s sequence. P_**ij**_ is less than 1 for all pairs of amino acids i,j used in the formula.

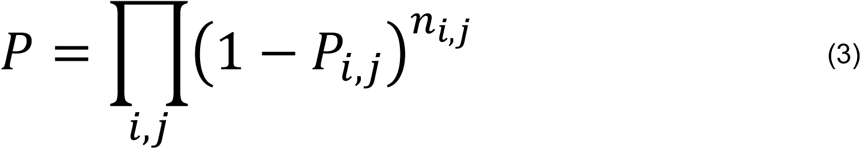

This P-value may detect stop codon avoidance (SCA) caused only by synonymous mutations. SCA due to functionally similar amino acid substitutions, like leucine to valine or isoleucine substitution in the case of −1 AORF frame, are not taken into account. There are methods that take into account such substitutions [41], however the groups of similar amino acids are defined for proteins from different organisms. In one organism any amino acid substitution can lead to the protein dysfunction.

Some genes (coding sequences) are only predicted by programs and their protein product existence is not proved experimentally or by similarity to reliable proteins. In the case of ORFs pair located on complementary strands with long overlap annotation programs typically choose one of these ORFs as putative coding sequence, and the choice might be wrong [42]. That is why we have produced symmetric calculations considering AORF as a coding sequence.

Obtained P-value distributions tend to uniform or have the peak for low P-values (S1 Fig.). This suggests for the relevancy of our P-value calculation method.

### Ka/Ks ratio calculations

We have used KaKs_Calculator 2.0 [43] to calculate Ka/Ks ratio for pairwise alignments using Yang and Nielsen method [44]. If Ka/Ks is calculated for the sequence in multiple alignment the average Ka/Ks by pairwise alignments is taken.

Applying this model to a pair of an essential gene and a non-coding AORF in frame −1 or - 3, we could expect to detect negative selection for the gene and “positive selection” (actually there is no selection) for the AORF because most of synonymous mutations in the gene lead to non-synonymous in the AORF translation. If the AORF is a protein coding gene, we can expect stabilizing selection also affecting it. Thus, we could detect “neutral selection” (Ka/Ks ≈ 1), or something remotely like this, for both AOGs in frame −1 or −3. The model presented in [38] is applicable only for cases of both strands encoding proteins. Our work was aimed to detect such cases but not to reveal the real type of selection affecting AOGs under “neutral selection” – the model [38] has not been used.

### Ka/Ks ratio calculations for genes

For each gene, we have found hits of its translation in 929 reference prokaryotic genomes using BLAST `tblastn` algorithm with the default parameters. Thus, a hit is an open reading frame with a translation similar to the translation of a gene. Genes Ka/Ks ratios have been calculated as average for pairwise alignments of a gene and hits taken from multiple alignment. No more than 30 best (one best per a genome) hits with the identity from 35% to 99% and longer than 180 bp have been used.

To confirm the patterns reported in [36], the same procedure has been applied for AORFs Ka/Ks ratio calculation.

### Alignments analysis

Multiple alignments have been constructed using MUSCLE with default parameters [45,46]. Phylogeny reconstruction has been done using FastME 2.07 [47] and `protdist` (The Dayhoff PAM matrix) from PHYLIP package (EMBOSS realization) [48]. ORFs translations BLAST [49] search has been done with `tblastn` (default parameters) against analyzed genomes and with `blastp` (word size 3) against NCBI RefSeq Proteins database. The search through HMM profiles has been conducted with HMMER3 package (http://www.hmmer.org).

The multiple alignment of functional sequences should contain domains and linkers between them. The regions corresponding to domains are expected to be more conservative than linkers. Therefore, conservative columns distribution should differ from uniform in alignments of functional sequences. Conservative columns uniformity in multiple alignments has been estimated with chi-squared test. A multiple alignment is split into windows of a particular size with a definite step. The number of conservative columns for window *i* (*c*_*i*_) and the mean 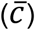 by all windows are calculated. *χ*^2^ statistics is calculated by formula 4. Thus, window size, step and conservation percent are the parameters.

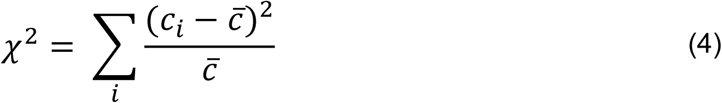

## Results

### AORFs lengths distribution

We have found 10 472 000 AORFs overlapping (overlap no less than 180 bp) with annotated genes in 929 analyzed prokaryotic genomes. On average it is about 10 000 long AORFs per one prokaryotic genome. A great part of these AORFs is unlikely to encode proteins.

The distributions of found AORFs lengths differ by AORFs frames (Fig. 1). Thus, features of genes influence stop codons frequency and their distribution along the antiparallel DNA strand. On the assumption of AORFs translation, synonymous mutations in a gene lead to the drastically different effect on AORFs amino acids sequences in different frames. In the case of frame −1 relatively gene there are six synonymous mutations in gene that are synonymous mutations in AORF: CAG ⇔ CAA in gene codon (Glu) are synonymous CTG ⇔ TTG in AORF (Leu); TCG ⇔ TCC (Ser) are CGA ⇔ GGA (Arg); CCT ⇔ CCG (Pro) are AGG ⇔ CGG (Arg) and symmetric ones, in which gene codons and AORF codons are exchanged. In the case of −3 frame there are no synonymous mutations in a gene that are synonymous in AORF. In the case of −2 frame third positions of codons on complementary strands are opposite to each other. Hence almost all synonymous mutations in third codon nucleotides are synonymous on the opposite DNA strand. Therefore AORFs in different frames should be analyzed separately.

**Fig. 1.**
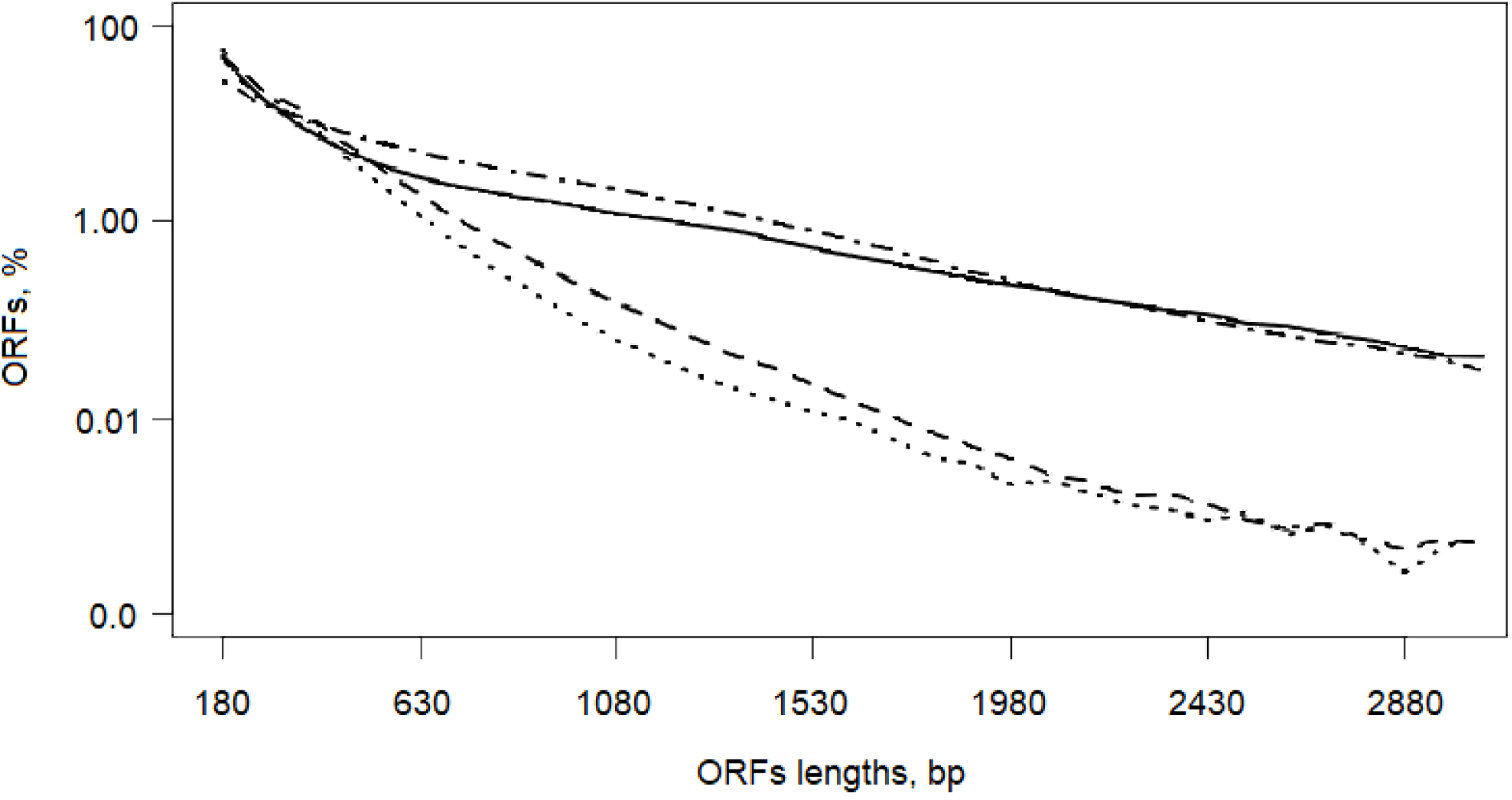
ORFs lengths distributions. Vertical axis is ORFs fraction of all ORFs in particular frame (logarithmic scale). Solid line – all ORFs in six reading frames; dot-dash line – AORFs in the frame −1; dashed line – AORFs in the frame −2; dotted line – AORFs in the frame - 3. The overlap with the annotated gene is not shorter than 180 bp for AORFs.

### AORFs with stop codon avoidance

Long AORFs in different frames significantly vary with respect to results of synonymous mutation in genes (Table 2). We have used P-value to estimate SCA caused by synonymous mutations (see Materials and Methods). All calculated P-values have been corrected with Benjamini-Hochberg correction at significance level 0.01. AORFs with reliable SCA are presented in S3 Table. If AORF actually encodes a protein, then both AORF and gene are in symmetrical relations. Indeed, if these AORFs with SCA are considered as genes and genes are considered as AORFs to them, then the probability of SCA has been calculated for genes.

**Table 2.** Long AORFs number in different frames.

| Frame | AORFs number | AORFs with SCA number | Pairs number with SCA on both strands<br>(% of AORFs with SCA) |
| --- | --- | --- | --- |
| -1 | 3 552 276 | 2 168 (0.061%) | 2 027 (93.5%) |
| -2 | 4 225 952 | 115 (0.003%) | 58 (50.4%) |
| -3 | 2 693 772 | 615 (0.023%) | 398 (64.7%) |
| Total | 10 472 000 | 2 898 (0.028%) | 2 483 (85.7%) |

Despite there are 39 (54 including stop codon-containing pairs) and 51 variants of amino acids residues pairs opposite to which stop codons of AORFs in frames −2 and −3 could appear, real frequencies of these pairs are low. As a result, we can see that AORFs with SCA are in the frame −1 in most cases. Furthermore, we do not take into account SCA due to the substitutions of amino acids residues in genes, though similar in function amino acids replacements like substitution of leucine to valine or isoleucine might contribute to the statistics of SCA in AORFs.

AORFs lengths distribution (S2 Fig.) is typical for prokaryotic genes in all 3 cases, i.e. peaks on the histograms are observed for ORFs lengths from 300 to 700 bp. Obtained statistics indicate that antiparallel overlapping genes in prokaryotic genomes with overlap longer than 180 bp might exist.

### AORFs to *dnaK* genes

Several cases of AORFs to *dnaK* genes in −1 frame were analyzed in earlier works [15-22]. That is why AORFs in the frame −1 opposite to *dnaK* genes have been examined for being protein-coding. We have found 978 orthologs of the *dnaK* gene from *E. coli* in 929 analyzed genomes (S4 Table). BLAST (`tblastn` with default parameters) hits with protein identity more than 45% and length more than 1500 bp (80% of *E.coli* DnaK protein) have been considered to be orthologous.

AORFs opposite to every *dnaK* ortholog in −1 frame have been found (S5 Table). Among them there are only 5 AORFs with reliable stop codon avoidance. Nevertheless, length histogram (Fig. 2) presents the peak for AORFs longer than 1800 bp. There are 282 such AORFs. All studied earlier AORFs to *dnaK* genes have approximately the same length. So these 282 AORFs have been chosen as candidates in protein-coding genes.

**Fig. 2.**
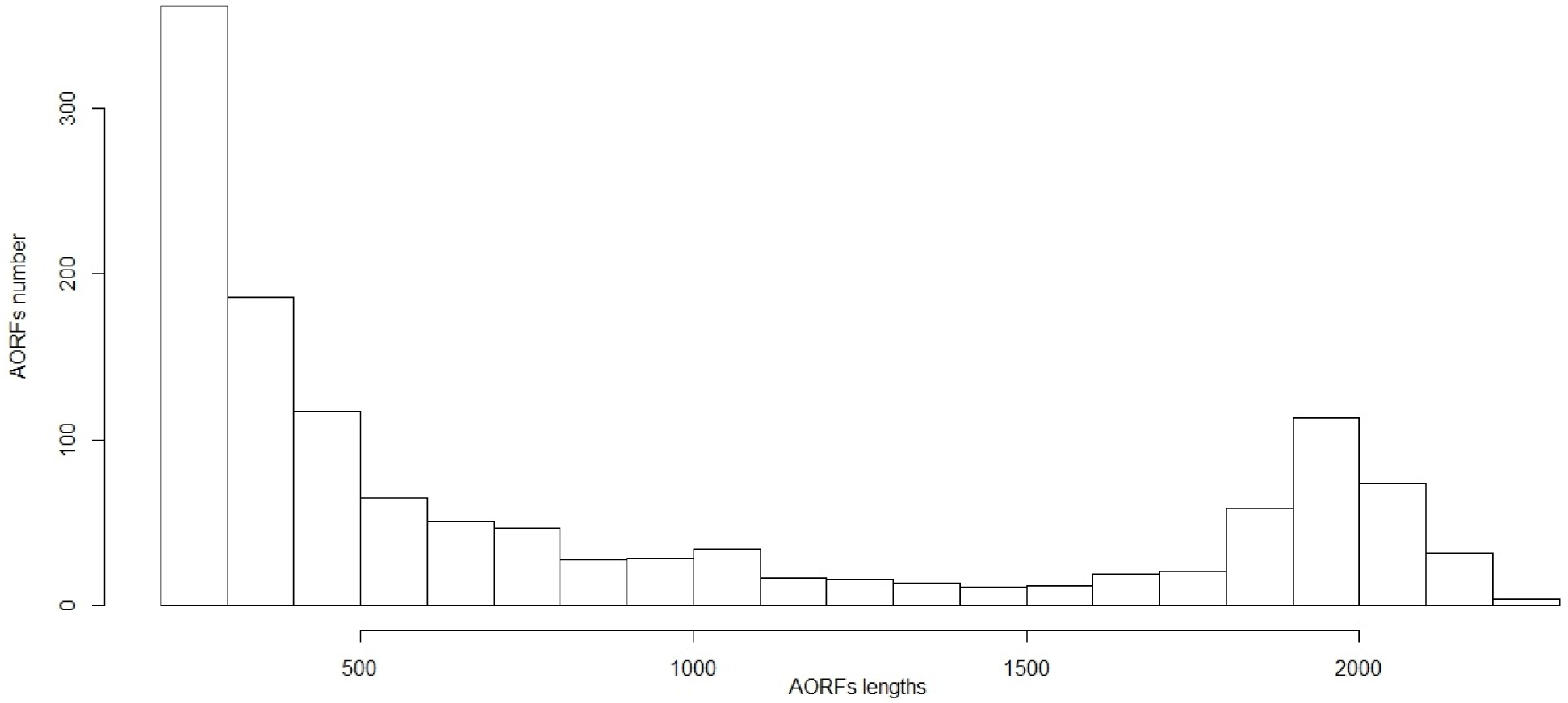
Lengths of AORFs to *dnaK* genes. Built by most long AORF in the frame −1 to obtained *dnaK* genes.

We have checked by HMM profile with HMMER3 (`hmmsearch` with default parameters) obtained from Pfam database that translations of all chosen AORFs belong to NAD-GH Pfam family PF10712 (S1 File).

Sequence alignments of enzyme families typically involve conserved motifs corresponding to the active center or other functional sites. We have constructed the multiple alignment of 282 AORFs translations longer than 600 amino acids residues using MUSCLE with default parameters (S2 File). Every chosen ORFs translations pair identity exceeds 30%. We have compared the distribution of conservative columns in long AORFs translation alignment and seed alignments of two other glutamate dehydrogenases Pfam families PF00208 and PF05088 (Table 3). AORFs alignment has been randomly rarefied to 100 sequences (comparable with seed alignments).

**Table 3.** Uniformity P-value (see methods) for alignments of long AORFs to *dnaK* genes, PF00208 and PF05088.

| Column conservation threshold | AORFs to *dnaK* rarefied alignment | PF00208 seed alignment | PF05088 seed alignment |
| --- | --- | --- | --- |
| 98% | 0.527 | 8.3e-05 | 0 |
| 100% | 0.677 | 1.15e-03 | 0 |

Window size 10 amino acids, window step 5 amino acids, peripheral columns with 95% of gaps have been removed.

Obtained results challenge the hypothesis that the long AORFs opposite to prokaryotic dnaK genes encode essential glutamate dehydrogenases. Our data show that absolutely conservative columns are distributed uniformly along translations of long AORFs to *dnaK* genes alignment but for two other families of glutamate dehydrogenases, we cannot say about uniformity (P-value < 0.05).

In addition, if long AORFs encode essential proteins (like glutamate dehydrogenases), then some homologs of this AORFS could occur without dnaK gene or pseudogene on the complement strand because of dnaK gene duplication and loss of its function in one copy. We have found no examples of such scenarios. Indeed, all BLAST (`tblastn` with default parameters) hits of these 282 AORFs translations with E-value less than 0.0001 are located opposite to *dnaK* homologs. All members of NAD-GH Pfam family are also located opposite to *dnaK* genes homologs. Similar results were reported in [24]. Reliable homology to tRNA synthetases proposed in several studies [26–28] has not also been found.

If long AORFs were encoding essential genes, we could see the common ancestor of *dnaK* genes and these AORFs pairs. Phylogenetic tree of DnaK proteins built with FastME (distances between DnaK proteins have been calculated with `protdist` from PHYLIP), however, shows that there is no clade on the tree in which all organisms would have long antiparallel to *dnaK* gene ORF and in other clades would not (S3 File).

To compare the character of selection affecting putative AORFs translation and DnaK we have used Ka/Ks ratio calculation. The phyla Actinobacteria and Proteobacteria have been chosen because there is the comparable number of genomes in our dataset and the comparable number of genomes with long AORFs located opposite to *dnaK* genes. Ka/Ks have been calculated for three multiple alignments of sequences: long AORFs, *dnaK* genes without long AORFs and *dnaK* genes with long AORFs (S6 Table). Obtained data demonstrate that AORFs are not under any type of selection while dnaK genes are under strong negative selection in spite of long AORF absence or presence (Fig. 3).

**Fig. 3.**
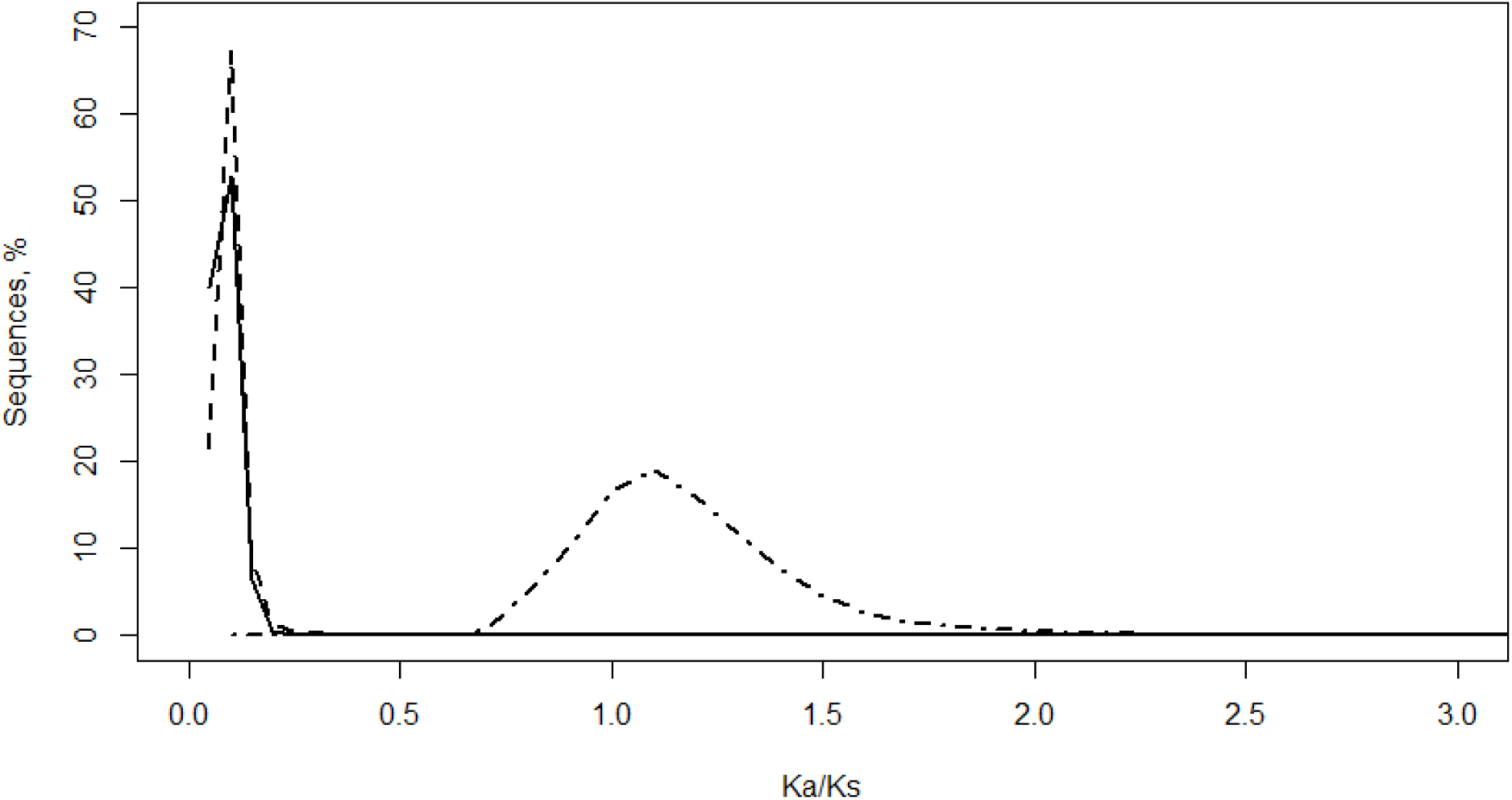
Ka/Ks ratio of *dnaK* genes with long AORFs (solid line), *dnaK* genes without long AORFs (dashed line) and long AORFs to *dnaK* genes (dot-dash line).

Thus, antiparallel to *dnaK* genes ORFs are unlikely to encode glutamate dehydrogenases which must be under stabilizing selection like a great part of essential genes (Fig. 5).

**Fig. 4.**
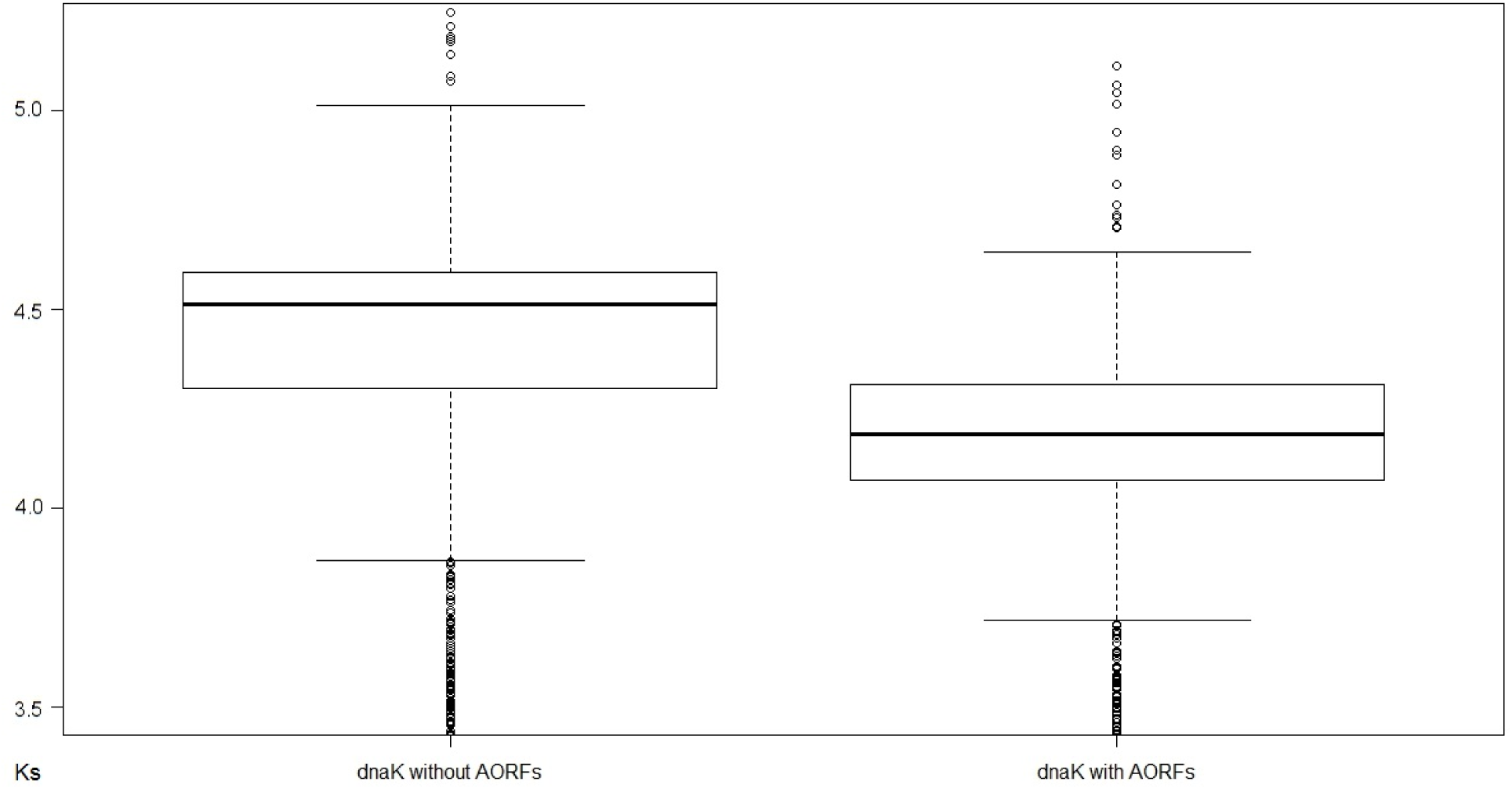
Ks value of *dnaK* genes without AORFs and with AORFs.

**Fig. 5.**
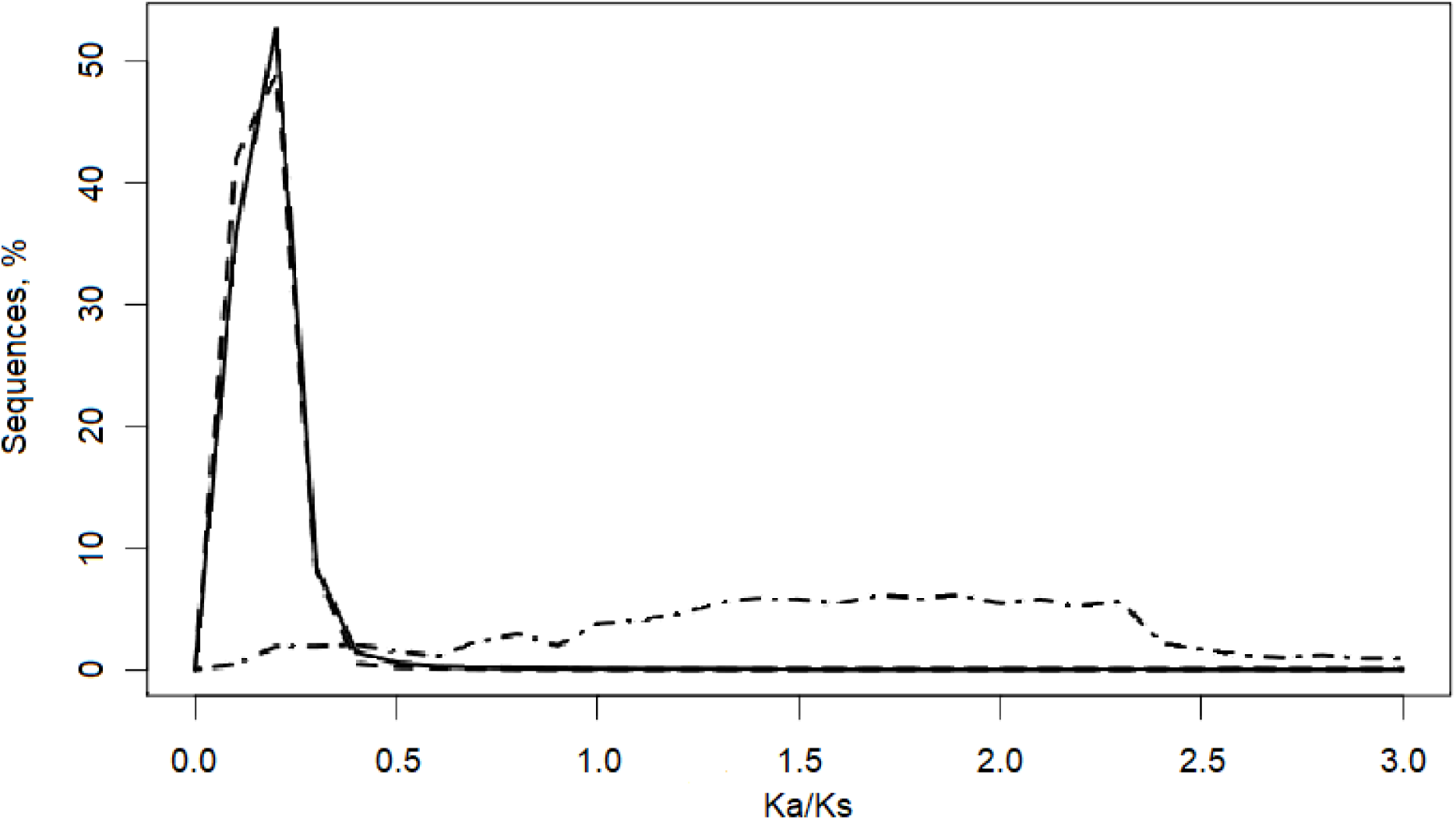
Ka/Ks ratio of genes with long AORFs with SCA (solid line), genes without long AORFs (dashed line), AORFs with SCA (dot-dash line).

On the other hand, Ks values (the evaluation of synonymous mutations rate) of *dnaK* genes without long AORFs are bigger (Fig. 4) than the same values calculated for *dnaK* genes with long AORFs (Mann-Whitney-Wilcoxon test P-value < 10-16). This difference indicates that statistically long AORFs lead to certain resistance to synonymous mutations in *dnaK* genes.

Similar analysis shows that *adm* gene, found opposite *dmdR1* gene in *S. coelicolor* A3(2) [12], is also unlikely to be essential for two reasons (the same procedures as for *dnaK* genes case have been used): first, a *dmdR1* homolog has been found opposite each *adm* homolog but there are *dmdR1* homologs without *adm* homologs on the reverse strand; second, phylogenetic tree reconstruction of *dmdR1* genes translations has not revealed the common ancestor of this pair (S4 File). Ka/Ks ratio values (Ka/Ks ≪ 1) obtained for *adm* genes are typical for genes under negative selection (S7 Table), however, these values can be observed because *adm* is in −2 frame relatively *dmdR1* that is under negative selection [36].

### AORFs to all genes. Ka/Ks ratio

We have calculated Ka/Ks ration for all annotated genes from analyzed genomes without long AORFs, all annotated genes with long AORFs (overlap longer than 180 bp), only for genes with long AORFs with SCA and for all long AORFs with SCA. More than 40% of long AORFs with SCA have no homologs (no reliable `tblastn` hits with identity from 35% to 99% and longer than 180 bp). We believe that these AORFs hardly encode essential proteins. Ka/Ks ratio has been calculated for the remaining 60% of AORFs.

Ka/Ks ratio calculations (Fig. 5) show that AORFs with SCA, if they are supposed encoding proteins, are under “positive selection”. These AORFs are not likely to be under any type of selection – the selection affecting functional genes sequences may lead to Ka/Ks values significantly differing from 1.0 for AORFs. Thereby, virtually all analyzed AORFs, as well as antiparallel to *dnaK* genes ORFs, seem to be not coding essential genes.

Distributions of Ka/Ks ratio for AORFs in different frames (S3 Fig.) demonstrate the specificity of the frame −2, which is presumably caused by coincidence of third positions in codons of genes and AORFs [36]. Indeed, synonymous mutations in a gene mostly are synonymous mutations in AORFs. Conversely, mutations in first and second positions of AORF codons correspond to mutations in second and first positions of a gene, which mostly are non-synonymous. Thus, low values of Ka/Ks ratio for AORFs can be explained by peculiarities of mutations in genes rather than AORFs amino acid conservation.

Interestingly, Ks value of genes with AORFs is smaller (Mann-Whitney-Wilcoxon test P-value = 0) than Ks of genes without AORFs (Fig. 6). This fact may indicate that a fraction of AORFs resist to synonymous changes in genes opposite to which they are located.

**Fig. 6.**
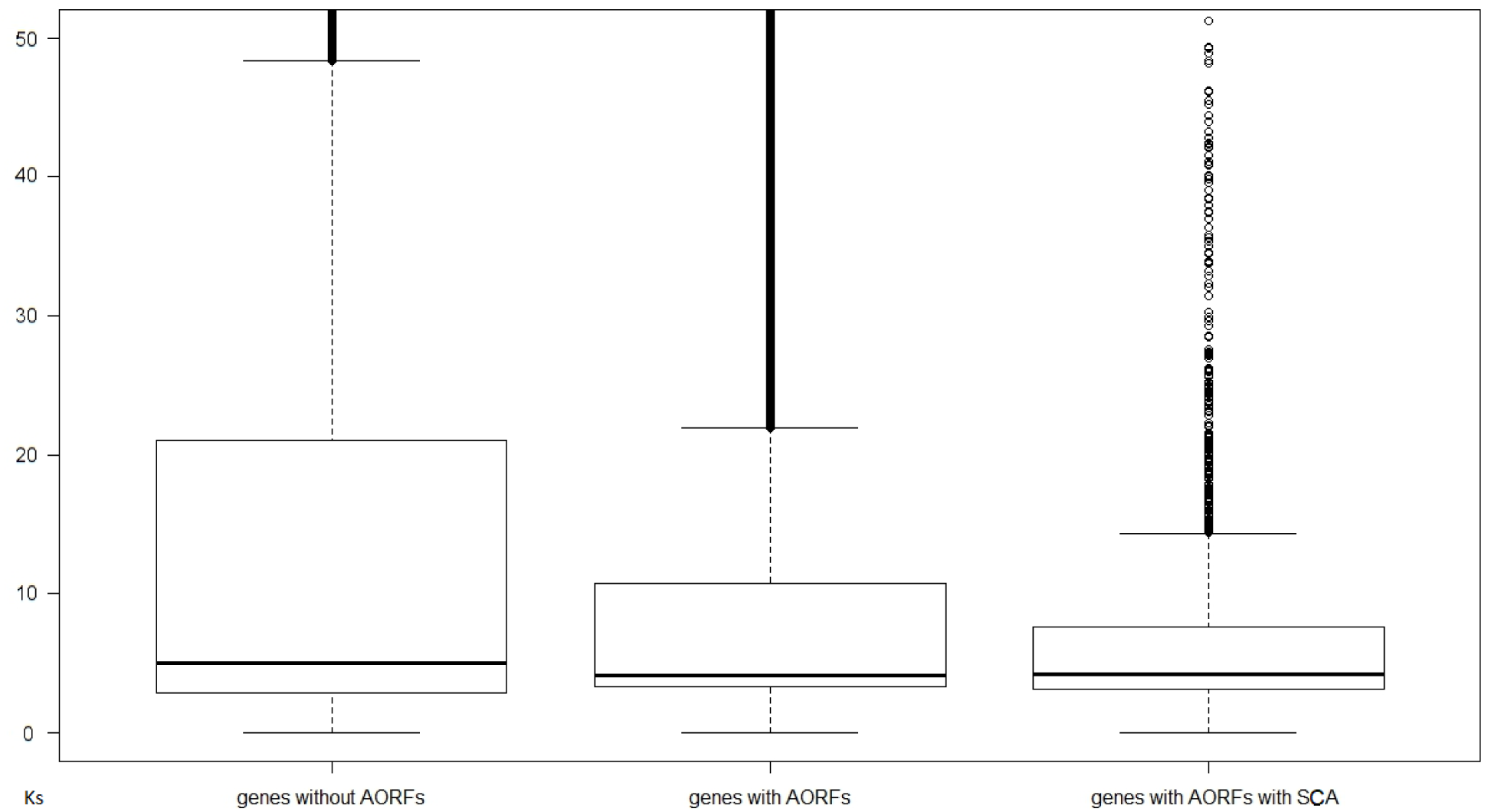
Ks rate of genes without AORFs, genes with AORFs and genes with AORFs for which stop codon avoidance (SCA) has been reliably detected.

It is worth noting that Ks values for genes with AORFs for which stop codon avoidance has been reliably detected in average less than Ks values for all genes with long AORFs. This result suggests that our method for AORFs P-value calculation could be used for AORFs functionality prediction should AORFs encode proteins.

### Candidates to AOGs

Comparing histograms of Ka/Ks for AORFs and genes (Fig. 5) we have chosen 0.4 as a threshold: genes and AORFs with Ka/Ks < 0.4 are considered to be under negative selection (S8 Table). The major part of (AORF, gene) pairs consist of genes under negative selection and AORFs that are not under any type of selection or their translations do not have homologs (Table 4). There are 2531 (1619 + 912) such pair. Presumably, genes are correctly annotated and these 2531 AORFs are unlikely to encode essential proteins.

**Table 4.** Classification of pairs (AORFs with stop codon avoidance (SCA), genes opposite AORFs) according to their evolutionary trends.

| **Genes**<br>**AORFs with SCA** | **No**<br>**homologs** | **Under**<br>**positive**<br>**selection** | **Under**<br>**negative**<br>**selection** | **Total** |
| --- | --- | --- | --- | --- |
| **No homologs** | 178 | 22 | 919 | 1119 |
| **Under “positive selection”** | 34 | 29 | 1619 | 1682 |
| **Under “negative selection”** | 6 | 14 | 77 | 97 |
| **Total** | 218 | 65 | 2615 | 2898 |

We have found 20 (6 + 14) pairs that could be the cases of erroneous coding strand choice by annotation programs. Ka/Ks ratios for AORFs are typical for genes under negative selection and Ka/Ks ratios for annotated genes are typical for positive selection or annotated genes do not have homologs at all. There are 8 pairs in which AORFs may encode proteins whereas gene annotations are false (S8 Table). Functions of the AORFs have been predicted with BLAST (`blastp` against NCBI RefSeq Proteins, word size 3).

There are 178 pairs such that neither gene translations nor AORF translations got any homolog. In addition, 85 (22 + 34 + 29) pairs of AORFs and genes are either under “positive selection” or their translations have no homologs. Although it is probable that in these cases neither genes nor AORFs encode essential proteins, some ORFs from these pairs might be real genes under positive selection.

We have revealed 77 pairs (among them 35 are in frame −2) in which both gene and AORFs translations can be under negative selection. They have been considered as the most probable candidates for antiparallel essential protein-encoding genes. If so, we may expect stop codon avoidance both in AORF and gene. There are 62 (S3 Table) such pairs (significance level 0.01 with Benjamini-Hochberg correction). Nevertheless, only one pair (AORF NC_014810|:1059006-1059758 and gene NC_014810|:c1059766-10583) might be AOGs (Table S9). AORF is in the frame −2, however, we have found its homolog without ORF on the reverse strand that is in favor of its functionality. The function of this AORF has not been found with BLAST `blastp` against NCBI RefSeq Proteins database (word size 3, E-value < 0.001). The gene translation is homologous to fumarate hydratase class II (WP_013469520).

It is worth noting that “interesting” values of Ka/Ks ratio for some pairs (Ka/Ks < 0.4 for a gene and its AORF) may be the result of average sensitivity to outliers.

## Discussion

The origin and evolution of antiparallel overlapping genes are of special interest. Many AOGs are presented in viral genomes. However, in prokaryotic genomes, only a few examples of antiparallel protein-coding genes with long overlaps were reliably confirmed. In this work, we investigated the abundance of this unusual phenomenon.

Our results, like published earlier estimations [9,31], show that the number of long AORFs is reliably bigger than expected according to the statistical model. Among 929 prokaryotic genomes 2898 AORFs might be considered as candidates in antiparallel overlapping genes because stop codons in their sequences are absent, presumably, not by chance and the lengths of their translations are typical for prokaryotic proteins: in most cases they are from 70 to 350 amino acids residues with mode at 250 amino acids residues (S2 Fig.). Such a big number may be explained either by the inaccuracy of statistical model or by a lot of AOGs existence in prokaryotic genomes.

The results presented in this work testify against AORFs encoding essential proteins, i.e. conservative proteins that have important functions for organism fitness. In the assumption that AORFs encode proteins, Ka/Ks ratio analysis shows that majority of AORFs with stop codon avoidance translations is not under any selection while genes opposite to which they are located are under strong negative selection. In case of AORFs in −2 frame relatively to genes, “negative selection” for AORFs translations must be explained by stabilizing selection for genes. It is known that a very small fraction of essential genes changing their functions is subjected to positive selection [37]. That is why AORFs are unlikely to encode essential proteins. The number of polypeptides like *adm* gene translation [12] might be much bigger than it is supposed.

We have characterized AORFs to *dnaK* genes in particulars. Data in favor these AORFs encoding glutamate dehydrogenases were published earlier [15–17,23,28]. And yet, their results are contradictory [25]. We have found 282 AORFs longer than 1800 bp (more than 600 amino acids residues in translation) opposite to *dnaK* genes (Fig. 2). All these AORFs are in −1 frame relatively to *dnaK* genes. Several arguments testify against the hypothesis that these AORFs encode essential proteins. First, like in all cohort of long AORFs, we have observed the prevalence of non-synonymous substitutions in AORFs to *dnaK* genes in compare with synonymous (Fig. 3). This observation is in agreement with the high frequency of synonymous mutations in *dnaK* genes. Indeed, practically every synonymous mutation of *dnaK* leads to non-synonymous substitution in AORFs codon due to −1 frame. At the same time, the assumption that protein-coding genes on the complement strand are not under any selection or under positive selection is unlikely to be the truth.

Second, phylogenetic analysis of DnaK proteins shows that AORFs are presented in many evolutionary independent branches. It is not typical for essential proteins.

Third, all homologs of AORFs to *dnaK* genes translations are located on the opposite strand to genes of DnaK homologs. For antiparallel overlapping gene encoding essential protein the loss of relations with ‘parent’ gene after duplication looks like believable scenario. In one copy *dnaK* might retain its function while in other copy antiparallel gene encoding protein could freely adopt its sequence breaking *dnaK* which became not essential. Nevertheless, we have found no protein such that its gene is located not opposite to *dnaK* even in NAD-GH Pfam family of glutamate dehydrogenases.

Forth, the alignment of long antiparallel to *dnaK* gene ORFs translations does not contain any conservative motifs typical for essential proteins – conservative positions are distributed more or less uniformly through the full alignment.

On the other hand, certain data testify in favor of long AORFs translation. First, experimental studies detecting translations of AORFs were published [2,11,12,14–17]. Second, obtained statistics indicate stop codon avoidance in some AORFs. Third, our data show that the frequency of synonymous substitutions in genes with long AORFs is less than in the case if long AORFs are absent (Fig. 6). This observation may be due to several limitations on substitutions in genes because of complement strand translation. Lower frequencies of synonymous mutations in some genes can also be caused by only frequent codons usage to maintain high expression level. Stop codons of an AORF in frame −1, for example, are located opposite rare codons for leucine and serine in the gene. There are AORFs located opposite constantly expressed genes: *dnaA, dnaK, rpoB*, etc. Thus, the occurrence of AORFs can be explained with codon usage optimization for some genes.

The analysis proposed by us can be used for genes annotation correction. If predicted only by annotation program gene has long AORF, the case of high Ka/Ks ratio value for gene translation and low for AORF translation looks like AORF is the true gene. Furthermore, should functional AORFs really exist, our method for P-value calculation could predict them. In general low P-value for an AORF indicates synonymous mutation rate restriction for the gene.

The deletion of AORFs was shown to change organisms phenotype but not to cause the death [2,11–13]. The effect of genes knockout was estimated in conditions typical for wild type strains. Actually, organisms do not have redundant genes. Thus, any gene must be useful for an organism in its natural conditions. However, the conditions changes can lead to some genes becoming harmful. This scenario is probable for genes studied in [2,11–13]. AORFs amount in several genomes is large enough to get a lot of various AORFs sets. Moreover, the lengths of AORFs in different strains of the same species can be significantly different. For example, in most *E. coli* strains including *E. coli* IAI39 AORFs to *dnaK* genes have lengths about 2000 bp while in *E. coli* K12 and its closest relatives these AORFs are shorter than about 1.5 times. Due to these observations, we do not exclude that several AORFs might contribute to the determination of some individual features of an organism, for example, such as size or color.

Summarizing all pros and cons, we speculate that several long AORFs are translated into polypeptides which do not have any highly specialized function and are not essential proteins. Nonetheless, they may play a role in a cell because of which stop codon avoidance and, possibly, conservation of particular amino acids residues are maintained.

## Acknowledgments

The work was supported by RSF grant 16-14-10319

## Supporting information

**S1 Fig. AORFs P-values distribution.**

**S2 Fig. AORFs with SCA lengths distribution.**

**S3 Fig. Ka/Ks ratio of AORFs with SCA in frame −1, frame −2 and frame −3.**

**S1 Table. 929 reference prokaryotic genomes.**

**S2 Table. Stop codons probabilities opposite to dipeptides.**

**S3 Table. AORFs with stop codon avoidance (SCA) and Ka/Ks ratios.**

**S4 Table. Orthologs of DnaK from *E. coli.***

**S5 Table. Antiparallel to *dnaK* genes ORFs in −1 frame.**

**S6 Table. Ka/Ks ratios of antiparallel to *dnaK* genes ORFs longer than 1800 bp, of *dnaK* genes without AORFs longer than 1800 bp, *dnaK* genes with AORFs longer than 1800 bp.**

**S7 Table. Ka/Ks ratios of *adm* genes translations, Ka/Ks ratios of *dmdR1* genes translations. S8 Table. BLAST hits for AORFs that are candidates to essential proteins coding genes.**

**S1 File. HMMER3 (`hmmsearch`) output by NAD-GH Pfam family HMM profile for translations of long antiparallel to *dnaK* genes ORFs.**

**S2 File. The alignment of antiparallel to *dnaK* genes ORFs translations longer than 600 amino acids.**

**S3 File. Phylogenetic tree built for 978 orthologs of DnaK from *E. coli*.** Orthologs opposite to which AORFs longer than 1800 bp are located are marked as “LONG AORF”.

**S4 File. Phylogenetic tree built for 252 *dmdR1* genes translations of Actinobacteria.**

## References

1. Neme R, Tautz D. Phylogenetic patterns of emergence of new genes support a model of frequent de novo evolution. BMC Genomics. 2013;14(1):1.

2. Fellner L, Simon S, Scherling C, Witting M, Schober S, Polte C, et al. Evidence for the recent origin of a bacterial protein-coding, overlapping orphan gene by evolutionary overprinting. BMC Evol Biol [Internet]. 2015 Dec [cited 2016 Sep 12];15(1). Available from: http://www.biomedcentral.com/1471-2148/15/283

3. Chirico N, Vianelli A, Belshaw R. Why genes overlap in viruses. Proc R Soc B Biol Sci. 2010 Dec 22;277(1701):3809–17.

4. Barrell BG, Air GM, Hutchison CA. Overlapping genes in bacteriophage phiX174. Nature. 1976 Nov 4;264(5581):34–41.

5. Normark S, Bergstrom S, Edlund T, Grundstrom T, Jaurin B, Lindberg FP, et al. Overlapping genes. Annu Rev Genet. 1983;17(1):499–525.

6. Lamb RA, Horvath CM. Diversity of coding strategies in influenza viruses. Trends Genet TIG. 1991 Aug;7(8):261–6.

7. Samuel CE. Polycistronic animal virus mRNAs. Prog Nucleic Acid Res Mol Biol. 1989;37:127–53.

8. Callen BP, Shearwin KE, Egan JB. Transcriptional interference between convergent promoters caused by elongation over the promoter. Mol Cell. 2004;14(5):647–656.

9. Pallejà A, Harrington ED, Bork P. Large gene overlaps in prokaryotic genomes: result of functional constraints or mispredictions? BMC Genomics. 2008;9(1):335.

10. Delaye L, DeLuna A, Lazcano A, Becerra A. The origin of a novel gene through overprinting in Escherichia coli. BMC Evol Biol. 2008;8(1):31.

11. Fellner L, Bechtel N, Witting MA, Simon S, Schmitt-Kopplin P, Keim D, et al. Phenotype of *htgA (mbiA*), a recently evolved orphan gene of *Escherichia coli* and *Shigella*, completely overlapping in antisense to *yaaW*. FEMS Microbiol Lett. 2014 Jan;350(1):57–64.

12. Tunca S, Barreiro C, Coque J-JR, Martín JF. Two overlapping antiparallel genes encoding the iron regulator DmdR1 and the Adm proteins control sidephore and antibiotic biosynthesis in Streptomyces coelicolor A3(2): Overlapping antiparallel genes and iron regulation. FEBS J. 2009 Sep;276(17):4814–27.

13. Silby MW, Levy SB. Overlapping protein-encoding genes in Pseudomonas fluorescens Pf0-1. PLoS Genet. 2008 Jun 13;4(6):e1000094.

14. Kim W, Silby MW, Purvine SO, Nicoll JS, Hixson KK, Monroe M, et al. Proteomic Detection of Non-Annotated Protein-Coding Genes in Pseudomonas fluorescens Pf0-1. Kelso J, editor. PLoS ONE. 2009 Dec 24;4(12):e8455.

15. Yang B, LeJohn HB. NADP (+)-activable, NAD (+)-specific glutamate dehydrogenase. Purification and immunological analysis. J Biol Chem. 1994;269(6):4506–4512.

16. LeJohn HB, Cameron LE, Yang B, MacBeath G, Barker DS, Williams SA. Cloning and analysis of a constitutive heat shock (cognate) protein 70 gene inducible by L-glutamine. J Biol Chem. 1994;269(6):4513–4522.

17. LeJohn HB, Cameron LE, Yang B, Rennie SL. Molecular characterization of an NAD-specific glutamate dehydrogenase gene inducible by L-glutamine. Antisense gene pair arrangement with L-glutamine-inducible heat shock 70-like protein gene. J Biol Chem. 1994;269(6):4523–4531.

18. Tavaria M, Gabriele T, Kola I, Anderson RL. A hitchhiker’s guide to the human Hsp70 family. Cell Stress Chaperones. 1996 Apr;1(1):23–8.

19. Morano KA. New tricks for an old dog: the evolving world of Hsp70. Ann N Y Acad Sci. 2007 Oct;1113:1–14.

20. Konstantopoulou I, Nikolaidis N, Scouras ZG. The hsp70 locus of Drosophila auraria (montium subgroup) is single and contains copies in a conserved arrangement. Chromosoma. 1998 Dec;107(8):577–86.

21. Monnerjahn C, Techel D, Mohamed SA, Rensing L. A non-stop antisense reading frame in the grp78 gene of Neurospora crassa is homologous to the Achlya klebsiana NAD-gdh gene but is not being transcribed. FEMS Microbiol Lett. 2000 Feb 15;183(2):307–12.

22. Rother KI, Clay OK, Bourquin JP, Silke J, Schaffner W. Long non-stop reading frames on the antisense strand of heat shock protein 70 genes and prion protein (PrP) genes are conserved between species. Biol Chem. 1997 Dec;378(12):1521–30.

23. Walker ND, McEwan NR, Wallace RJ. Overlapping sequences with high homology to functional proteins coexist on complementary strands of DNA in the rumen bacterium Prevotella albensis. Biochem Biophys Res Commun. 1999 Sep 16;263(1):58–62.

24. Finn RD, Coggill P, Eberhardt RY, Eddy SR, Mistry J, Mitchell AL, et al. The Pfam protein families database: towards a more sustainable future. Nucleic Acids Res. 2016 Jan 4;44(D1):D279–285.

25. Williams TA, Wolfe KH, Fares MA. No Rosetta Stone for a Sense-Antisense Origin of Aminoacyl tRNA Synthetase Classes. Mol Biol Evol. 2009 Feb 1;26(2):445–50.

26. Rodin SN, Ohno S. Two types of aminoacyl-tRNA synthetases could be originally encoded by complementary strands of the same nucleic acid. Orig Life Evol Biosphere J Int Soc Study Orig Life. 1995 Dec;25(6):565–89.

27. Carter CW, Duax WL. Did tRNA synthetase classes arise on opposite strands of the same gene? Mol Cell. 2002 Oct;10(4):705–8.

28. Rodin AS, Rodin SN, Carter CW. On Primordial Sense–Antisense Coding. J Mol Evol. 2009 Nov;69(5):555–67.

29. Chandrasekaran SN, Yardimci GG, Erdogan O, Roach J, Carter CW. Statistical evaluation of the Rodin-Ohno hypothesis: sense/antisense coding of ancestral class I and II aminoacyl-tRNA synthetases. Mol Biol Evol. 2013 Jul;30(7):1588–604.

30. Merino E, Balbás P, Puente JL, Bolívar F. Antisense overlapping open reading frames in genes from bacteria to humans. Nucleic Acids Res. 1994 May 25;22(10):1903–8.

31. Mir K, Neuhaus K, Scherer S, Bossert M, Schober S. Predicting Statistical Properties of Open Reading Frames in Bacterial Genomes. Battista JR, editor. PLoS ONE. 2012 Sep 24;7(9):e45103.

32. Miyata T, Yasunaga T. Evolution of overlapping genes. Nature. 1978 Apr 6;272(5653):532–5.

33. Pavesi A, De Iaco B, Granero MI, Porati A. On the informational content of overlapping genes in prokaryotic and eukaryotic viruses. J Mol Evol. 1997 Jun;44(6):625–31.

34. Krakauer DC. Stability and evolution of overlapping genes. Evol Int J Org Evol. 2000 Jun;54(3):731–9.

35. Rogozin IB, Spiridonov AN, Sorokin AV, Wolf YI, Jordan IK, Tatusov RL, et al. Purifying and directional selection in overlapping prokaryotic genes. Trends Genet TIG. 2002 May;18(5):228–32.

36. Mir K, Schober S. Selection Pressure in Alternative Reading Frames. White BA, editor. PLoS ONE. 2014 Oct 1;9(10):e108768.

37. Firth AE, Brown CM. Detecting overlapping coding sequences with pairwise alignments. Bioinforma Oxf Engl. 2005 Feb 1;21(3):282–92.

38. Sabath N, Landan G, Graur D. A Method for the Simultaneous Estimation of Selection Intensities in Overlapping Genes. Pybus OG, editor. PLoS ONE. 2008 Dec 22;3(12):e3996.

39. Sabath N, Graur D. Detection of Functional Overlapping Genes: Simulation and Case Studies. J Mol Evol. 2010 Oct;71(4):308–16.

40. Zhang R. DEG: a database of essential genes. Nucleic Acids Res. 2004 Jan 1;32(90001):271D–272.

41. Lèbre S, Gascuel O. The combinatorics of overlapping genes. J Theor Biol. 2017 21;415:90–101.

42. Poptsova MS, Gogarten JP. Using comparative genome analysis to identify problems in annotated microbial genomes. Microbiology. 2010 Jul 1;156(7):1909–17.

43. Wang D, Zhang Y, Zhang Z, Zhu J, Yu J. KaKs_Calculator 2.0: a toolkit incorporating gamma-series methods and sliding window strategies. Genomics Proteomics Bioinformatics. 2010 Mar;8(1):77–80.

44. Yang Z, Nielsen R. Estimating synonymous and nonsynonymous substitution rates under realistic evolutionary models. Mol Biol Evol. 2000;17(1):32–43.

45. Edgar RC. MUSCLE: multiple sequence alignment with high accuracy and high throughput. Nucleic Acids Res. 2004;32(5):1792–7.

46. Edgar RC. MUSCLE: a multiple sequence alignment method with reduced time and space complexity. BMC Bioinformatics. 2004 Aug 19;5:113.

47. Lefort V, Desper R, Gascuel O. FastME 2.0: A Comprehensive, Accurate, and Fast Distance-Based Phylogeny Inference Program. Mol Biol Evol. 2015 Oct;32(10):2798–800.

48. Rice P, Longden I, Bleasby A. EMBOSS: the European Molecular Biology Open Software Suite. Trends Genet TIG. 2000 Jun;16(6):276–7.

49. Altschul SF, Gish W, Miller W, Myers EW, Lipman DJ. Basic local alignment search tool. J Mol Biol. 1990 Oct 5;215(3):403–10.

